# cudaMMC - GPU-enhanced Multiscale Monte Carlo Chromatin 3D Modelling

**DOI:** 10.1101/2023.06.12.544609

**Authors:** Michał Wlasnowolski, Paweł Grabowski, Damian Roszczyk, Krzysztof Kaczmarski, Dariusz Plewczynski

**Author notes:** equally contributed (Michał Wlasnowolski, Paweł Grabowski, Damian Roszczyk).

## Abstract

**Motivation:** Investigating the 3D structure of chromatin provides new insights into transcriptional regulation. With the advancements in 3C new-sequencing techniques such as ChiA-PET and Hi-C, there has been a substantial increase in the volume of collected data, necessitating faster algorithms for chromatin spatial modelling. This study presents the cudaMMC method, which utilises the Simulated Annealing Monte Carlo approach extended by GPU-accelerated computing to generate ensembles of chromatin 3D structures efficiently.

**Results:** The *cudaMMC* calculations demonstrate significantly faster performance and lower (better) model scores compared to our previous method on the same workstation. *cudaMMC* substantially reduces the computation time required for generating ensembles of large chromatin models, making it an invaluable tool for studying chromatin spatial conformation.

**Availability:** Open-source software and manual and sample data are freely available on https://github.com/SFGLab/cudaMMC

## 1 Introduction

In recent years, the development of high-throughput sequencing methods and chromosome conformation capture (3C) technology has shown the significant influence of chromatin spatial conformation on genetic expression. Dynamic changes in the 3D structure at various levels of DNA spatial organisation: from the entire chromosomes, chromosomal territories, domains (TAD, CCDs), or single chromatin loops, affect the transcription level of individual genes (reviewed in (Chiliński *et al*., 2021)). There is plenty of evidence that rearrangements of the spatial chromatin structure could alter gene expression. These changes may occur dynamically in the cell environment induced by heat stress or cell differentiation (Ray *et al*., 2019, Pei *et al*., 2020), as well as caused by genetic mutations, viruses infections, Structural Variations and DNA methylations (Sadowski *et al*., 2019, Lazniewski *et al*., 2019, Heinz *et al*. 2019). To investigate this phenomena, various algorithms for generating chromatin 3D structures were developed (Kadlof *et al*., 2020, Di Pierro *et al*., 2016), among others one is the *3D-GNOME* approach (Szałaj *et al*., 2016). *3D-GNOME* allows for generating ensembles of the 3D models of DNA using the simulated annealing Monte Carlo approach based on a map of chromatin contacts mediated by specific proteins like CTCF or RNAPII that play a crucial role in chromatin spatial organisation. This modelling technique considers the DNA hierarchical structure organisation, starting from chromosome positioning through chromatin domains (TADs, CCDs) to a single chromatin loop shape. Due to the substantial development of 3C techniques, the data volumes of the chromatin contacts have increased significantly. This substantially increases the computation time needed to model the ensemble of large chromosomal structures, making it less useful for genome-wide modelling. To address this issue, we developed cudaMMC, which extended 3D-GNOME by implementing parallel acceleration using GPUs, resulting in a significant speed-up of calculations while maintaining modelling quality. This is necessary for calculation of spatial distance distribution between specific genomic loci, for which purpose we applied cudaMMC recently in the 3D-GNOME web server update (Wlasnowolski *et al*. 2020) for statistical analysis of changes of distances between gene promoters and enhancers. We also added a new option for the output file format, mmCIF, which enables the presentation and analysis of chromatin 3D models using commonly used molecular 3D viewers, such as UCSF Chimera (Pettersen et al., 2004).

## 2 Materials and methods

### 2.1 *3D-GNOME -* CPU-oriented approach

The *3D-GNOME* method consists of two stages. In the first one, a chromosome is divided into several regions based on PET cluster interaction patterns so that each region can correspond to one topological domain. Next, Monte Carlo simulated annealing is used to position beads to minimise energy function, considering the distance between beads corresponding to different regions. The second stage models the position of chromatin anchors within each domain independently based on energy terms. Next, chromatin loops are modelled by inserting sub-anchor beads between adjacent anchors, wherefore their position and shape are set using minimised energy function again.

### 2.2 cudaMMC approach GPU-extended

A successful GPU algorithm must find a balance between local and global operations to minimise synchronisation and maximise parallel computations. GPU processing is based on so-called warps: synchronised cooperative groups of 32 threads, which may perform partially independent steps and exchange data using intrinsic commands or highly specialised shared memory. Our implementation of parallel simulated annealing based on the Monte Carlo method uses warps to perform random moving of the beads (Fig. 1*A*). A single warp optimises a single bead using 32 threads to try random moves of that bead iteratively. If some position is better, it is stored for further evaluation. Finally, a warp selects a minimum from its threads using parallel reduction. This strategy allows for the parallel fitting of thousands of beads in 32 different directions each.

**Fig 1.**
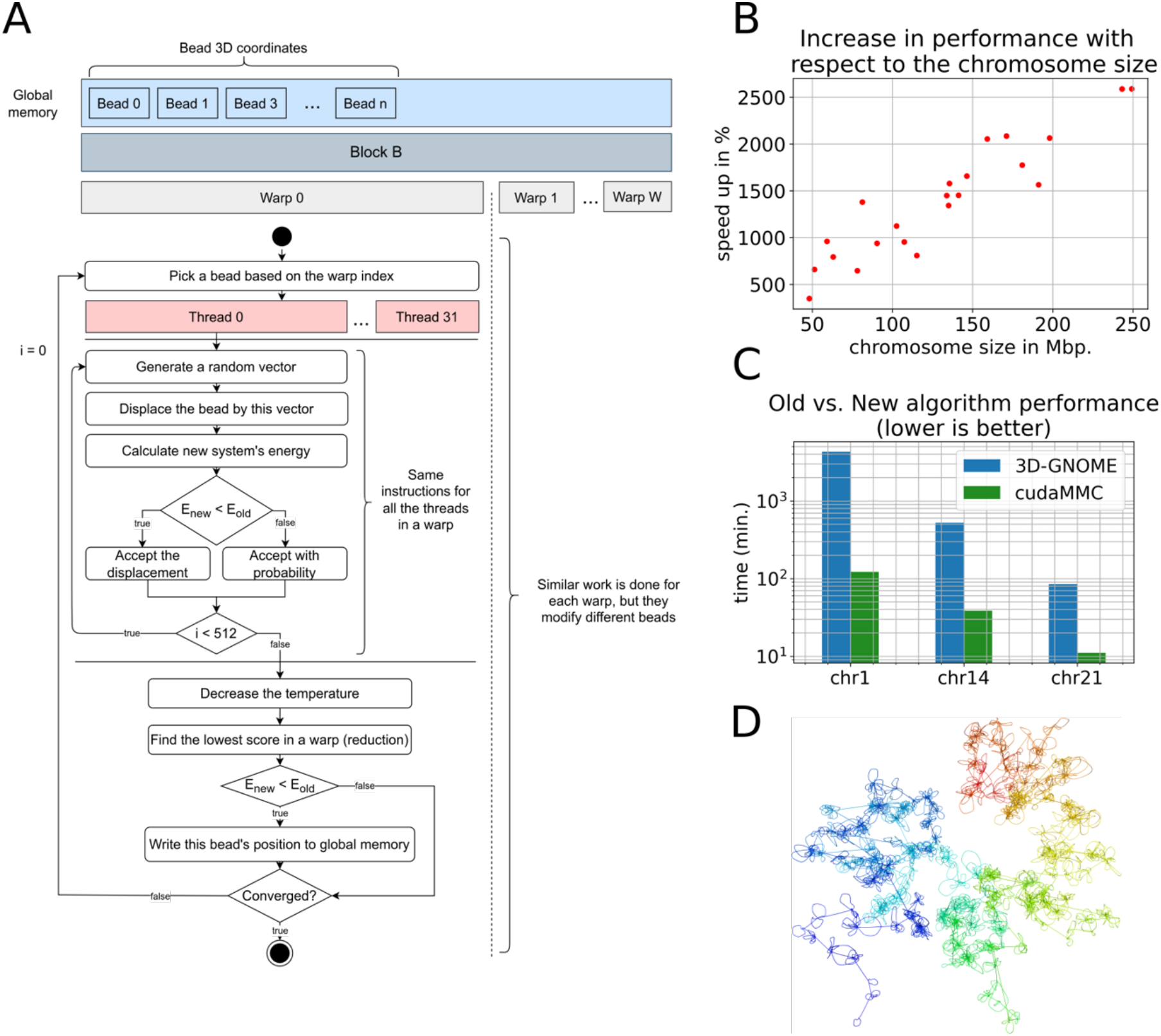
(A) Pipeline of the Parallel Simulated Annealing CUDA-Based Algorithm for Chromatin 3D Structure Modeling. (B) Speedup of the parallel algorithm with respect to the chromosome size. (C) Comparison of the performance between the 3D-GNOME (old) and cudaMMC (new) algorithms based on generating ensembles of 100 models for selected chromosomes. (D) Full chromosomal chromatin 3D structure of the chr1 based on CTCF ChIA-PET chromatin interactions for the GM12878 cell line mapped on GRCh37.

### 2.3 Chromatin interaction data

We compared *cudaMMC* and *3D-GNOME* algorithms on *long-read ChIA-PET* CTCF chromatin interaction data for the GM12878 cell line mapped on GRCh37, for which data *3D-GNOME* was designed. We have also performed modelling tests using *cudaMMC* on *in situ ChIA-PET* data for the GM12878 cell line mapped on GRCh38. We omitted the *3D-GNOME* performance testing on GRCh38 data because of the excessive computational time required caused by the massive increase of data gained by the new *ChIA-PET* method.

## 3 Comparison of performance between cudaMMC and 3D-GNOME

We compared the performance of the cudaMMC and 3D-GNOME methods on a workstation with a NVIDIA Pascal GPU architecture and an Intel Core i9-7920X CPU. The comparison was based on data sizes of varying sizes. The cudaMMC algorithm resulted in a speed-up of 3x to 25x for a single chromosome modelling, depending on the chromosome size (Figure 1B). The biggest advantage of the algorithm speed-up was observed for generating ensembles of models. Using GRCh37 data, we performed ensembles consisting of 100 models each using both methods. The cudaMMC algorithm decreased computation time from ∼85 to ∼11 min. for chr21, from ∼8.5h to ∼38 min. for chr14, and from ∼3 days to 2h for chr1. Moreover, we tested the performance of the cudaMMC algorithm for whole chromosomal modelling based on in situ ChIA-PET data mapped on GRCh38. The ensemble of 100 models generated for chr1 took ∼8h, for chr14 ∼2h 20 min., and for chr21 ∼21 min. (Figure 1C). These results were obtained on Pascal architecture, but cudaMMC might also be configured on Turing or Amper NVIDIA devices perhaps with even better results. We added a tool for converting output into mmCIF format to make it more accessible for common usage. We chose this format instead of PDB because it can represent a higher number of beads for one structure, which is necessary for high-resolution whole chromosome models (Figure 1D). Overall, our results show that the cudaMMC algorithm can significantly improve performance for 3D modelling of chromatin structures, making it a valuable tool for researchers in this field.

## 4 Conclusions

In the cudaMMC method, parallelising calculations on the GPU enabled a massive reduction in computation time compared to the previous CPU-oriented approach of 3D-GNOME. This speed-up in calculations enables the generation of a high number of model ensembles in a reasonable time frame, thereby offering an opportunity for statistical analysis. For example, as part of the update to the 3D-GNOME web server, we have recently added a tool for statistical analysis of ensembles, which calculates changes in distances between gene promoters and enhancers of two model ensembles that differ in chromatin loop patterns caused by Structure Variants. To facilitate this, we have set up the cudaMMC software on the NVIDIA DGX A100 cluster, enabling the running of multiple modelling tasks simultaneously on several GPU cards and facilitating analysis based on new, large datasets of spatial chromatin data. We believe that cudaMMC, the GPU-accelerated 3D-GNOME modelling engine, will significantly enhance scientists’ investigations into various aspects of chromatin 3D structures.

